# Leaf rolling as indicator of water stress in *Cistus incanus* from different provenances

**DOI:** 10.1101/131508

**Authors:** Giacomo Puglielli, Loretta Gratani, Laura Varone

## Abstract

The relationship between leaf rolling and physiological traits under imposed water stress conditions was analyzed in *C. incanus* representative saplings collected at different altitudes (i.e. Castelporziano, 41°45′N, 12°26′E, 0 m a.s.l. and Natural Park of Monti Lucretili, 42°33′N, 12°54′E, 750 m a.s.l) and grown *ex-situ.* The hypothesis that leaf rolling reflected physiological changes occurring during water stress irrespective to the different acclimation to cope with water stress was tested.

On the whole, the results show that leaf rolling is associated to an increased sub-stomatal CO_2_ concentration (*C*_i_) and a decreased carboxylation efficiency (*C*_e_). Moreover, leaf rolling in *C. incanus* leaves might be involved in protecting the PSII complex under water stress during the progressive inhibition of photosynthetic metabolism.

## Introduction

Leaf movements are common adaptive responses to stress factors in plants (Kadioglu et al. 2012). Leaf movement affects physiological performance because of the influence of orientation on leaf energy balance (Gamon and Pearcy 1989). Various reports (Ehleringer and Forseth 1980; Forseth and Ehleringer 1982; Gamon and Pearcy 1989; Mooney and Ehleringer 1978) show that diaheliotropism maximizes carbon gain by increasing incident photosynthetic photon flux density (PPFD) or minimizes incident radiation, resulting in more favorable leaf temperatures and water status during drought. Moreover, according to Ludlow and Björkman (1984) paraheliotropism contributes to avoid leaf photoinhibition under drought stress. Among leaf movements leaf rolling is an hydronastic mechanism involved in plant responses to stress factors (Kadioglu et al. 2012) such as water stress (Kadioglu et al. 2012). There is evidence (Heckathorn and De Lucia, 1991; Kadioglu and Terzi 2007; Kadioglu et al. 2012; Nar et al. 2009) that under water stress conditions leaf rolling affects stomatal conductance and consequently photosynthesis. Nevertheless, how it occurs is not clear yet. Abd Allah (2009) highlighted that leaf rolling reduces leaf surface exposed to sun light energy causing stomata closure and limiting CO_2_ uptake. On the contrary, O’Toole and Cruz (1980) found that partial leaf rolling in leaves with adaxial stomata increased stomatal conductance by providing a more favorable microenvironment such as a higher relative humidity. However, the contribution of leaf rolling on stomatal conductance under water stress depends on several factors including stomatal distribution as well as the degree and pattern of stomatal opening at low leaf water potential (Heckathorn and DeLucia 1991).

In Mediterranean ecosystems, the distribution of the dominant growth form and habitus is related to water availability. According to a gradient of increasing aridity, there is a decrease in the transpiring surfaces up to the complete lack of leaves in drought deciduous shrubs, associated with drought-evading annual species (De Micco and Aronne 2009). An intermediate form between evergreen and drought deciduous species is represented by seasonally dimorphic species. Unlike drought deciduous plants, in seasonally dimorphic species the decrease in transpiring surfaces during the drought period is not complete.

In particular, to cope with drought stress these species develop twigs with short internodes (brachyblasts) characterized by small xeromorphic leaves in summer, and twigs with longer internodes (dolichoblasts) with larger mesomorphic leaves in winter (De Micco and Aronne 2009). Seasonal leaf dimorphism has been reported to be an adaptive strategy to the seasonal climatic changes occurring in Mediterranean habitats (Aronne and De Micco, 2001; Christodoulakis et al. 1990; Kyparissis et al., 1997; Orshan 1964). Moreover, in seasonal dimorphic Mediterranean species leaf rolling has been decribed as a mechanism to reduce light interception (Aronne and De Micco 2001; Gratani and Bombelli 1999).

Seasonal dimorphic *Cistus* sp. are known to avoid photochemistry drought induced impairment by leaf movements such as variation in leaf angle (Flexas et al. 2014; Werner et al. 1999, 2001) and leaf rolling. Among these species *Cistus incanus* L. is a typical Mediterranean shrub species distributed along the coastal belt of the Central-Eastern Mediterranean, Northern Africa and Western Asia, extending from sea level to 800 m a.s.l. (Pignatti 1982).

The aim of this research was to analyze the relationship between leaf rolling and physiological variables in two populations of *Cistus incanus* growing at the altitudinal limits of its distribution area and being subjected to different selective pressures. In particular, the population growing at the highest altitude faces lower chilling temperatures during winter and a higher amount of precipitation during the year associated to a reduced extent of summer drought. On the contrary, the costal population faces a reduced amount of precipitation during the year associated to a prolonged summer drought. The hypothesis that leaf rolling reflected physiological changes occurring during water stress irrespective to the different acclimation to cope with water stress was tested.

## Materials and methods

### Study site and plant material

The study was carried out in July 2014 at the experimental garden of Sapienza University of Rome (41°54′N, 12°31′E; 41 m a.s.l.). Three-year old saplings of *C. incanus* grown from seeds collected in June 2012 in the Mediterranean maquis developing along the *Latium* coast near Rome (Castelporziano, 41°45′N, 12°26′E, 0 m a.s.l.) and at the Natural Park of Monti Lucretili (42°33′N, 12°54′E, 750 m a.s.l.) were considered. Saplings (twenty saplings per provenance) were cultivated in plastic pots (32 cm diameter and 29 cm depth) containing silt (5-8 %), clay (16-39 %) and sand (56-75%) (pH 7.2 – 7.5) and grown inside a growth chamber under constant photosynthetically active radiation (PPFD) of 600 μmol m^-2^ s^-1^ (12 h), at 25/20 °C light/dark average temperature and 50/40% relative air humidity.

The provenance sites are characterized by a Mediterranean climate (Fig. 1). In particular, at Castelporziano the mean minimum air temperature (T_min_) of the coldest months (January and February) is 4.8 ± 2.3°C, the mean maximum air temperature (T_max_) of the hottest months (July and August) is 30.5 ± 1.8°C, and the yearly mean air temperature (T_mean_) is 16.4 ± 6.1°C. The dry period is from mid-May to August (48.3 mm of total rainfall in this period). Total annual rainfall equals 824 mm with the greatest part occurring in autumn and in winter (data from Meteorological Station of Roma Capocotta, SIARL, Arsial, for the period 2004–2013).

**Fig. 1.**
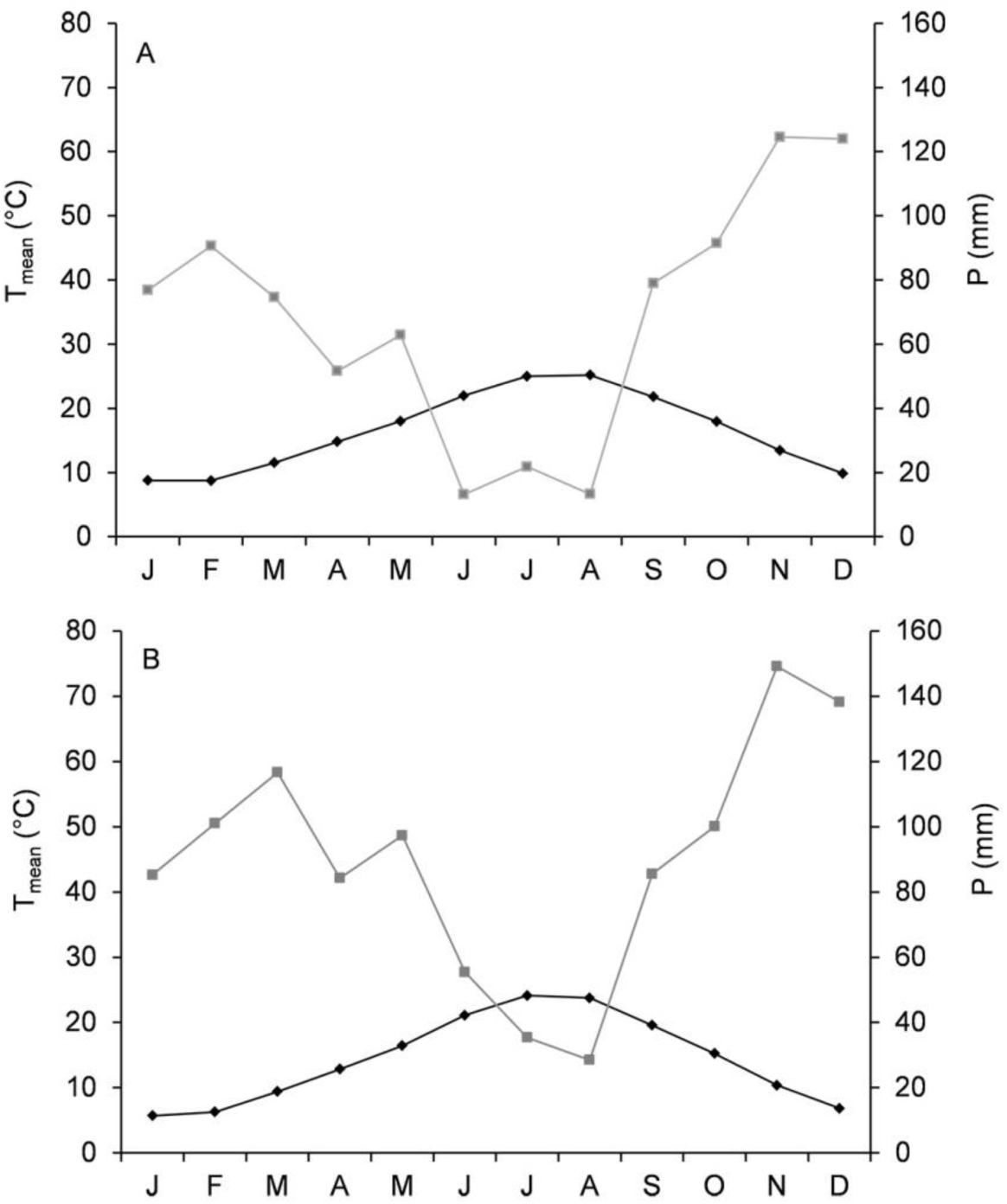
Bagnouls-Gaussen’s diagram (Time series 2004-2013) for A: Castelporziano 41°45′N, 12°26′E, 0 m a.s.l., and for B: the Natural Park of Monti Lucretili (42°33′N, 12°54′E, 750 m a.s.l.).

At the Natural Park of Monti Lucretili the mean minimum air temperature (T_min_) of the coldest months (January and February) is 0.5 ± 1.6°C, the mean maximum air temperature (T_max_) of the hottest months (July and August) is 32.7 ± 1.6°C, and the yearly mean air temperature (T_mean_) is 14.3 ± 6.7°C. The dry period is from mid-June to August (63.9 mm of total rainfall in this period). Total annual rainfall equals 1077 mm with the greatest part occurring in autumn and in winter (data from Meteorological Station of Palombara Sabina, SIARL, Arsial, for the period 2004–2013).

### Experimental procedure

Until the onset of the experiment (on July 5^th^), twenty saplings from Castelporziano and from Monti Lucretili (CP and LC saplings, respectively) were watered regularly to field capacity. Gas exchange, leaf water status, chlorophyll fluorescence and leaf rolling index measurements for the control saplings were performed on the first day of the experiment when all the saplings were well watered. Thereafter, the water stress was imposed by withholding water from ten CP and ten LC saplings, randomly arranged (stressed saplings, CPs and LCs, for Castelporziano and Monti Lucretili, respectively). In each sampling day the measurements were carried out on six randomly selected saplings per provenance.

The remaining ten saplings per provenance were kept under daily irrigation and measured to verify that the considered parameters maintained constant values through the experiment. The water stress experiment was stopped when stomatal conductance in CPs and LCs was below 0.05 mol m^-2^ s^-1^ indicative of a severe water stress condition (Medrano et al. 2002). On the whole the experiment lasted four days. Thereafter, the experimental days are indicated as D1 (first experimental day), D2 (two days after the beginning of the experiment) and D3 (four days after the beginning of the experiment).

### Gas exchange

During the experiment, maximum net CO_2_ assimilation rate (*A*_max_, μmol CO_2_ m^-2^ s^-1^), stomatal conductance (*g*_s_, mol H_2_O m^-2^ s^-1^), leaf transpiration (*E*, mmol H_2_O m^-2^ s^-1^) and substomatal CO_2_ concentration (*C*_i_, ppm) were measured with an open infrared gas analyser system (LCpro+, ADC, UK), equipped with a leaf chamber (PLC, ADC, UK).

Measurements were carried out while the natural inclination of the leaves was maintained. Between each measurement the IRGA was calibrated for CO_2_ and water vapor following the instructions of the manufacturers.

Six measurements on young fully expanded sun leaves per each selected control and stressed sapling were carried out every two days in the morning (11.00–12.30 h) at saturating PPFD (1500 μmol photon m^-2^ s^-1^) provided by the light source (LCpro+ Lamp unit). Before each measurement, the leaves were acclimated to saturated light conditions (*c*. 15?20 min). CO_2_ concentration in the leaf chamber (*C*_a_) was set at 400 μmol CO_2_ mol^-1^air, and relative air humidity of the incoming air ranged between 40% and 60%. Leaf temperature (*T*_l_, °C) was set at 25°C and varied by 1% during measurements. Apparent carboxylation efficiency (*C*_e_) was also determined by the ratio between A_max_ and *C*_i_ (Flexas et al. 2001).

### Chlorophyll fluorescence

Measurements of chlorophyll fluorescence were carried out on the same leaves of gas exchange measurements, using a portable modulated fluorometer (OS5p, Opti-Sciences, USA).

Chlorophyll fluorescence measurements were carried out at saturating PPFD (i.e. 1500 μmol photon m^-2^ s^-1^) ensuring a uniform light distribution on leaf surface while maintaining an inclination of the fluorometer pulse source at 45°.

The actual quantum efficiency of the photosystem II (Φ_PSII_) was calculated according to Genty et al. (1989) as:

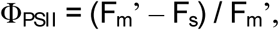

where F_m_’ was the maximum fluorescence obtained with a saturating pulse (*c*. 8,000 μmol m^-2^ s^-1^ PPFD) and F_s_ was the steady-state fluorescence of illuminated leaves (1,500 μmol m^-2^ s^-1^ PPFD). The rate of electron transport rate (ETR) was calculated, according to Krall and Edwards (1992) as:

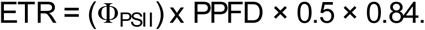

### Leaf water status

Leaf water potential (ψ_leaf_, MPa) was measured in control and stressed plants by a pressure bomb (SKPM 1400, Sky Instruments, Powys, UK).

The samples were enclosed in a bag previously saturated with CO_2_ and water vapor in order to avoid water losses from stomata.

Measurements were carried out in each sampling occasion on four leaves per each of the considered sapling after gas exchange measurements. In addition, relative water content (RWC, %) was measured as:

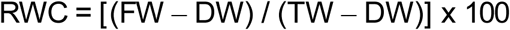

where FW was the leaf fresh weight, DW was the leaf dry weight after drying at 80 °C until constant weight was reached, and TW was the leaf weight after re-hydration until saturation for 48h at 5 °C in the darkness.

### Leaf rolling index

Leaf rolling index (LRI, %) was calculated on the same leaves used for gas exchange measurements according to Li et al. (2010) as:

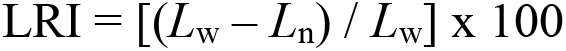

where *L*_w_ was the maximum leaf blade width and *L*_n_ the natural distance of the leaf blade margins. For LRI measurements *L*_w_ was measured only at D1 in order to avoid any confounding effect of leaf manipulation on gas exchange and chlorophyll fluorescence measurements.

### Statistical analysis

One-way ANOVA was performed to evaluate the differences between CPs and LCs and between stressed and control saplings at *p* ≤ 0.05. Multiple comparisons were analyzed by a Tukey test. Regression analysis was used to explore the relationships among the considered variables. Relationships were considered significant at *p* ≤ 0.01.

Kolmogorov–Smirnov and Levene tests were used to verify the assumptions of normality and homogeneity of variances, respectively. All data are shown as mean ± standard deviation (s.d.). All the statistic tests were performed by a statistical software package (Statistica.8, Stasoft, USA).

## Results

### Gas exchange

The daily gas exchange measurements are shown in and Figure 2.

**Fig. 2.**
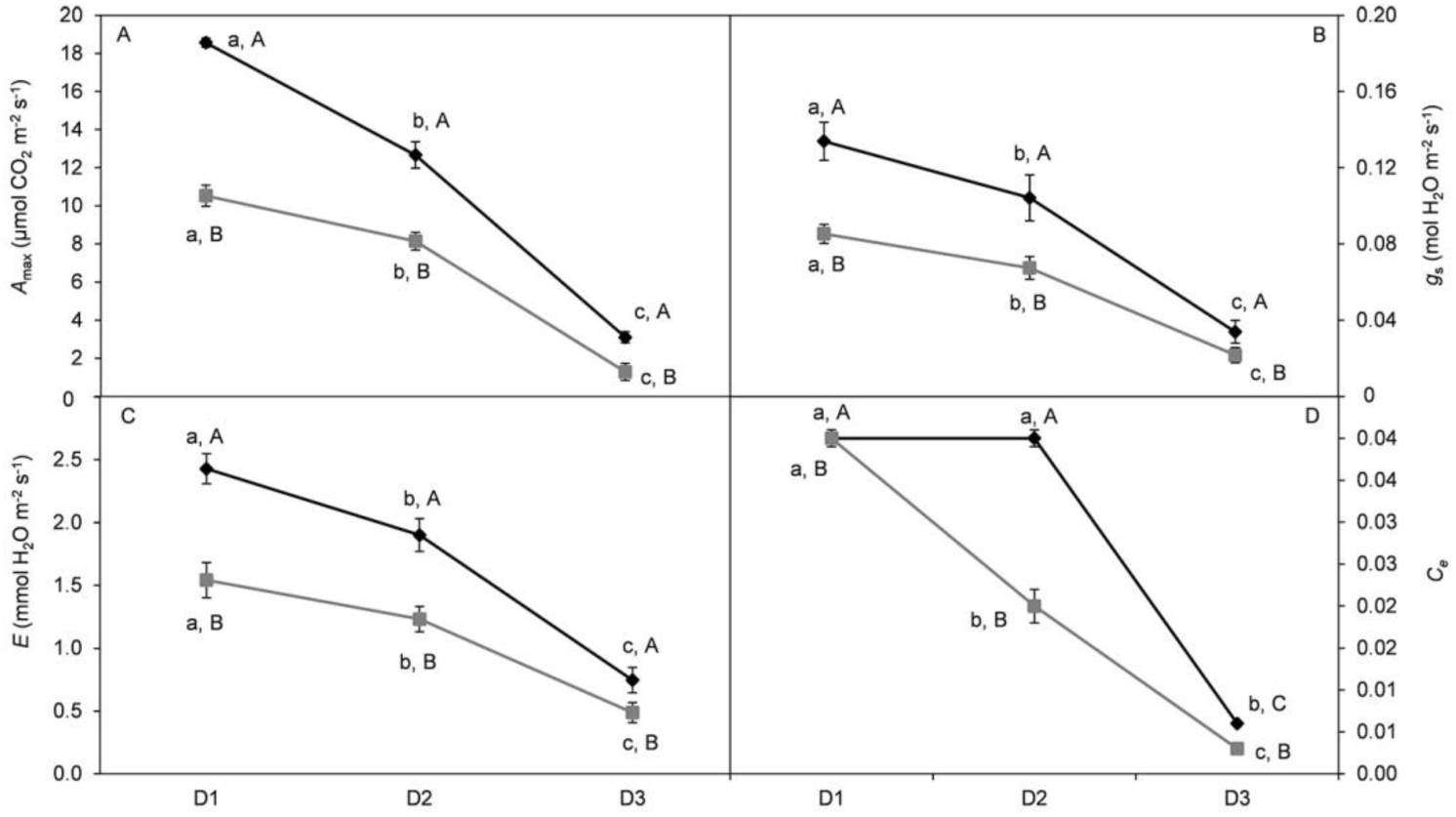
Trends of A: maximum net CO_2_ assimilation rate (*A*_max_), B: stomatal conductance (*g*_s_), C: leaf transpiration (*E*), and D: carboxylation efficiency (*C*_e_) measured during the first experimental day (D1), two days after the beginning of the experiment (D2) and four days after the beginning of the experiment (D3) of stressed *Cistus incanus* saplings from Castelporziano (CPs, black line) and from the Natural Park of Monti Lucretili (LCs, gray line). Mean values (± SE) are shown (n =36). Lowercase letters show significant differences between experimental days, capital letters show significant differences between CPs and LCs saplings at p = 0.05.

During the experiment the highest *A*_max_ values were measured at D1 for both CPs (18.6±2.6 μmol CO_2_ m^-2^ s^-1^) and LCs (10.5±2.6 μmol CO_2_ m^-2^ s^-1^). *A*_max_ decreased by 32% and 83%, at D2 and D3, respectively in CPs, and by 23% and 88%, respectively, in LCs (Fig. 2A). *g*_s_ showed the same *A*_max_trend dropping below 0.05 mol H_2_O m^-2^ s^-1^ at D3 for both CPs and LCs (Fig. 2B).

In CPs saplings *C*_i_ was 260±9.8 ppm at D1 decreasing by 6% at D2 and increasing by 9% at D3 compared to D1. *C*_i_ increased during the experiment by 11% and 41% at D2 and D3, respectively, compared to D1 (224±22 ppm). *C*_e_ did not vary between D1 and D2 in CPs (0.04±0.01) while it decreased by 75% at D3. In LCs, *C*_e_ decreased by 50% and 92%, at D2 and D3, respectively, compared to D1 (0.04±0.01) (Fig. 2D).

The control plants did not show any significant variation in gas exchange parameters during the experiment both in CPc and LCc.

### Chlorophyll fluorescence

The ETR and Φ_PSII_ values measured during the experiment are shown in Fig. 3.

**Fig. 3.**
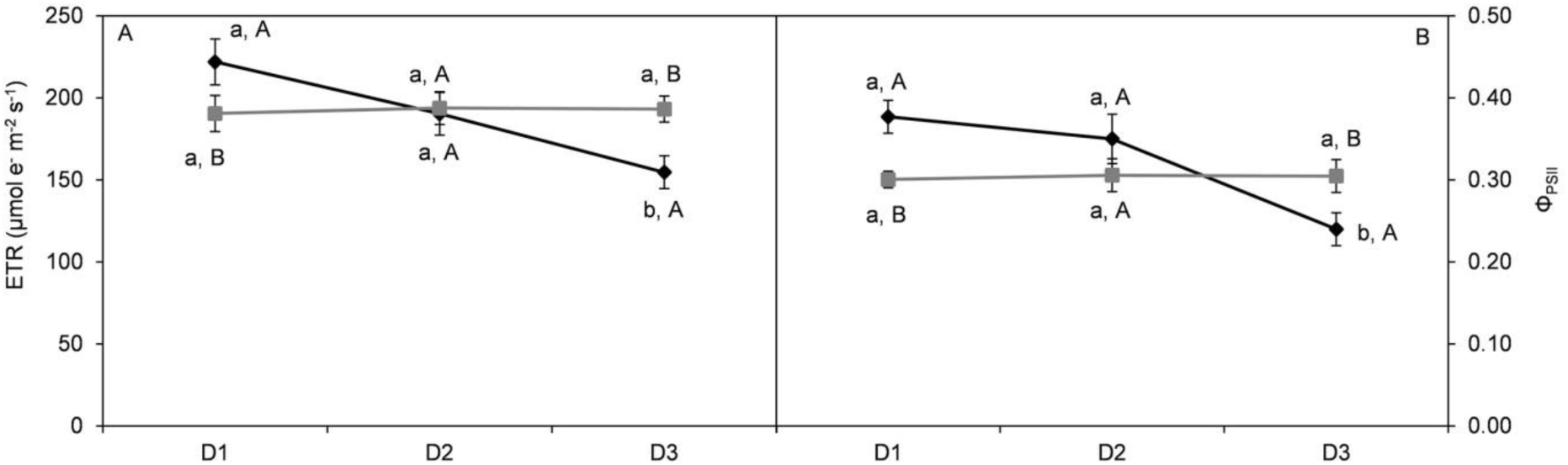
Trends of: A: rate of electron transport (ETR), and B: the actual quantum efficiency of the photosystem II (Φ_PSII_) measured during the first experimental day (D1), two days after the beginning of the experiment (D2) and four days after the beginning of the experiment (D3) of stressed *Cistus incanus* saplings from Castelporziano (CPs, black line) and from the Natural Park of Monti Lucretili (LCs, gray line). Mean values (± SE) are shown (n =36). Lowercase letters show significant differences between experimental days, capital letters show significant differences between CPs and LCs saplings at p = 0.05.

**Fig. 4.**
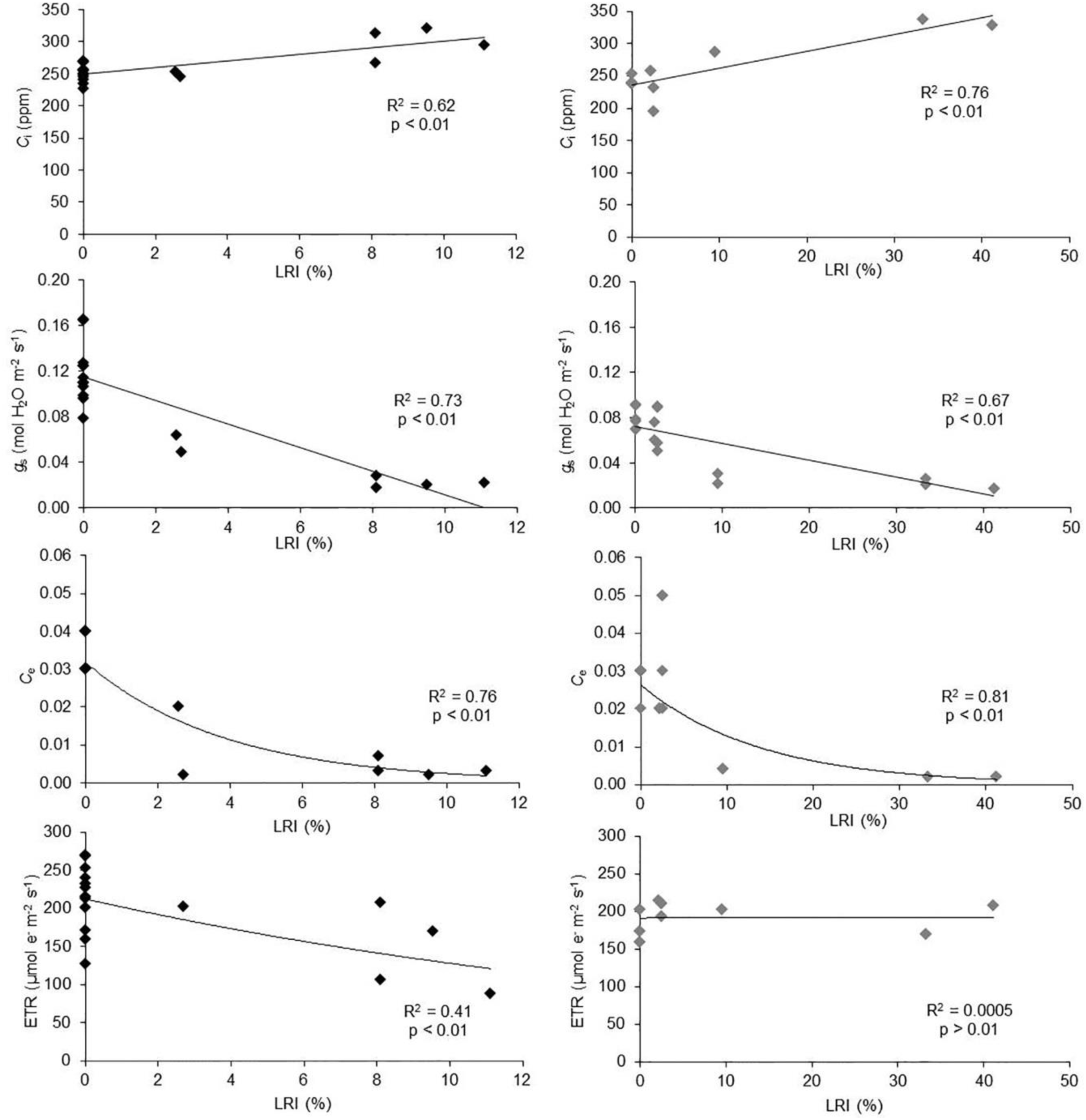
Relationships between leaf rolling index (LRI) and substomatal CO_2_ concentration (*C*_i_), stomatal conductance (*g*_s_), carboxylation efficiency (*C*_e_) and rate of electron transport (ETR) of stressed *Cistus incanus* saplings from Castelporziano (CPs, black dots, left column) and from the Natural Park of Monti Lucretili (LCs, gray dots, right column). Daily mean values per sapling were used as experimental units (*n* = 18, *p* ≤ 0.01).

In particular, in CPs ETR decreased by 14% and 30% at D2 and D3, respectively, compared to D1 (221.9±20.2 μmol e^-^ m^-2^ s^-1^) (Fig. 3A). Φ_PSII_ showed the same ETR trend with the highest value at D1 (0.38±0.05) decreasing by 8% and 37% at D2 and D3, respectively (Fig. 3B).

In LCs saplings ETR and Φ_PSII_ through the experiment did not vary significantly between the experimental days (Fig. 3A, B).

### Leaf water status

The measured values of RWC and ψ_leaf_ are shown in Table 1.

**Table 1.**
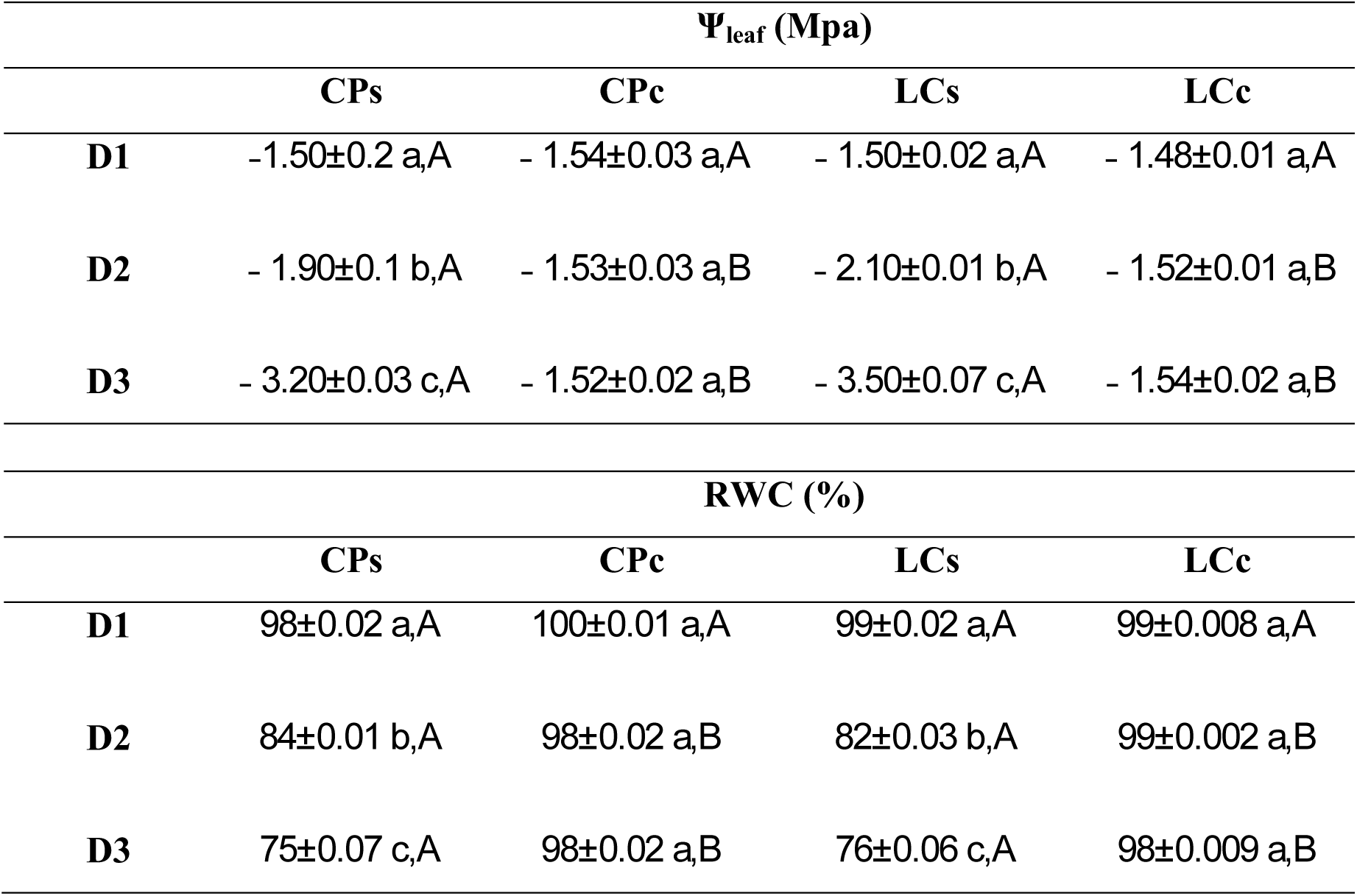
Leaf water potential (ψ_leaf_) and relative water content (RWC) measured during the first experimental day (D1), two days after the beginning of the experiment (D2) and four days after the beginning of the experiment (D3). CPs = stressed saplings of *Cistus incanus* from Castelporziano, CPc = control saplings of *Cistus incanus* from Castelporziano, LCs = stressed saplings of *Cistus incanus* from Natural Park of Monti Lucretili LCc = control saplings of *Cistus incanus* from Natural Park of Monti Lucretili. Mean values (± SE) are shown (n = 24). Lowercase letters show significant differences between experimental days, capital letters show significant differences between control and stressed saplings at p = 0.05.

CPs and LCs showed a significant decrease (by 113% and 133%) in ψ_leaf_ compared to D1 (- 1.50±0.2 and -1.50±0.02 MPa for CPc and LCc, respectively). RWC was 100% at D1 in both control and stressed saplings decreasing by 75% in both CPs and LCs at D3. Control plants did not show any significant variation of RWC and ψ_leaf_ during the experiment.

### Leaf rolling index

At D1 both CPs and LCs did not show any symptom of rolling. At D2, LRI did not increase in CPs while it slightly and significantly increased by 2% in LCs. LRI increased by 7% and 28% in CPs and LCs, respectively, at D3.

Control saplings did not show any symptom of rolling during the experiment (Table 2).

**Table 2.**
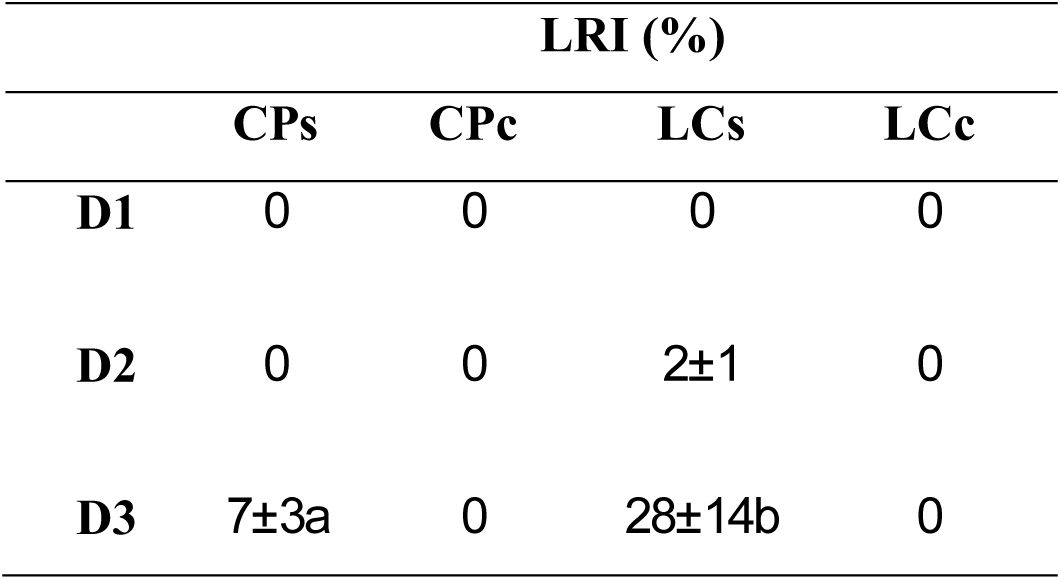
Leaf rolling index (LRI) measured during the first experimental day (D1), two days after the beginning of the experiment (D2) and four days after the beginning of the experiment (D3). CPs = stressed saplings of *Cistus incanus* from Castelporziano, CPc = control saplings of *Cistus incanus* from Castelporziano, LCs = stressed saplings of *Cistus incanus* from Natural Park of Monti Lucretili, LCc = control saplings of *Cistus incanus* from Natural Park of Monti Lucretili. Mean values (± SE) are shown (n = 24). Lowercase letters show significant differences between CPs and LCs saplings at p = 0.05.

LRI showed a linear relationship (p < 0.01) with *C*_i_ (R^2^=0.62 and R^2^=0.73, for CPs and LCs, respectively) and *g*_s_ (R^2^= -0.62 and R^2^= -0.73, for CPs and LCs, respectively). LRI was also significantly related with *C*_e_ in both CPs and LCs (R^2^= -0.76 and R^2^= -0.81, respectively) while it is significantly correlated with ETR only in CPs (R^2^= -0.41).

## Discussion

The results highlight a different response of CPs and LCs to water stress, which reflects their different provenances. In fact, LCs saplings, which experience a shorter period of drought in their natural environment in respect to CP saplings, are characterized by a higher sensitivity to water stress highlighted by a faster metabolic impairment.

In particular, both CPs and LCs showed the lowest values of ψ_leaf_ (-3.2±0.03 and -3.5±0.05 MPa, respectively) at D3 reaching a RWC value of 75.5±0.7%. When expressed as percentage of the control, *A*_max_, *g*_s_ and *E* trends were similar in CPs and LCs. Nevertheless, ETR and *C*_e_ were the parameters that showed the greatest differences between CPs and LCs.

According to Lawlor and Cornic (2002), the decrease in RWC increases *C*_i_. However, in CPs, *C*_i_ decreases at D2 (6%) and increased by 9% at D3, compared to D1. In LCs *C*_i_ increases through the experiment. In response to stomatal closure, CO_2_ inside the leaf initially declines then it increases as drought become more severe (Lawlor 1995). The decrease in *C*_i_ at D2 for CPs plants suggest that stomatal limitations dominate under moderate water stress, irrespective of any metabolic impairment (Flexas and Medrano 2002). However, at a certain stage of water stress *C*_i_ frequently increases highlighting the predominance of non-stomatal limitations of photosynthesis. Usually, the point at which *C*_i_ starts to increase occurs around *g*_s_ values of 0.05 mol H_2_O m^-2^ s^-1^ (Flexas and Medrano 2002). Accordingly, CPs show the highest value of *C*_i_ at D3 when *g*_s_ was 0.03 mol H_2_O m^-2^ s^-1^. On the contrary, LCs show the occurrence of non-stomatal limitations in D2 when *g*_s_ was 0.07 mol H_2_O m^-2^ s^-1^. The results are confirmed by the different *C*_e_ trend in CPs and LCs. In fact, CPs show *C*_e_ values equal to 100% of the control until D2, decreasing by 25% of the control at D3, while in LCs *C*_e_ decreases by 50% and 7.5% of the control at D2 and D3, respectively.

A RWC higher than 75% has not effect on photosynthetic metabolism (Lawlor and Cornic 2002). Nevertheless, the results of the experiment highlight a faster progressive inhibition of metabolism in LCs, associated to a higher RWC (i.e. 80%), than in CPs.

Water stress exposes plants to photo-inhibition by reducing PSII efficiency and ETR (Cabrera 2002). During the experiment both ETR and Φ_PSII_ decrease in CPs showing the lowest values at D3, while in LCs they do not vary significantly compared to the control. In particular, the ETR and Φ_PSII_ decrease in CPs at D2 may be interpreted as a down-regulation mechanism at lower *A*_max_ (Biehler and Fock 1996; Cornic and Massacci 1996; Haupt-Herting and Fock, 2000, 2002). Since the rate of electron transport at saturating photon flux is determined by sink capacity for electrons (such as photosynthesis at high RWC), a decreased sink capacity for electrons results in an increased non-photochemical energy dissipation (Lawlor and Cornic 2002). This may be justified by Φ_PSII_ reduction observed in CPs while the ETR decrease at D3 may be the result of non-stomatal limitations. On the contrary, the lack of variation in ETR and Φ_PSII_ in LCs suggests that the redox system under water stress is in a reduced state due to continued electron transport and absence of sinks (Lawlor and Cornic 2002), as confirmed by the significant decrease in *C*_e_ through the experiment. Moreover, the constant ETR and the *C*_i_ increase in LCs during the experiment could be also due to an increased photorespiration rate. In fact, decreasing RWC has long been known to increase the ratio of photorespiration to photosynthesis (Lawlor 1976; Lawlor and Fock 1975). Nevertheless, fluorescence and O_2_ isotope analysis showed that despite in stressed leaves the photorespiration to photosynthesis ratio increases, the absolute photorespiration rate was not so large like in unstressed leaves, so fewer electrons may be used (Biehler and Fock 1996).

During the experiment LCs show leaf rolling (i.e. LRI = 2±1%) at D2 increasing to 28±14% at D3, while in CPs leaf rolling (LRI = 7±3%) appears at D3. Since leaf rolling is a hydronastic mechanism the delayed leaf rolling in CPs indicates the ability to better sustain turgor, despite water stress, compared to LCs according to Kadioglu et al. (2012).

Moreover, LRI shows a significant (p < 0.01) linear relationship with *C*_i_ (R^2^=0.62 and R^2^=0.73, for CPs and LCs, respectively) and *g*_s_ (R^2^= -0.62 and R^2^= -0.73, for CPs and LCs, respectively), suggesting that LRI reflects the leaf physiological changing which occur during water stress. LRI is also significantly related with *C*_e_ in both CPs and LCs (R^2^= -0.76 and R^2^= -0.81, respectively) while it is significantly correlated with ETR only in CPs (R^2^= -0.41). The lack or the weakness of the relationship between LRI and ETR, associated to a strong negative relationships between LRI and *C*_e_ in both CPs and LCs, suggests that LRI is involved in protecting PSII under drought condition (Nar et al. 2009) during the progressive inhibition of photosynthetic metabolism.

In conclusion, the results highlighted that leaf rolling is related to the physiological variables in both CPs and LCs despite their different response to water stress. Thus, leaf rolling in seasonal dimorphic Medieterranean species may be used as morphological index not only to monitor the water status in the field but also to evaluate the progressive inhibition of photosynthetic metabolism irrespective to the different acclimation to cope with water stress.

